# Quantifying the genomic determinants of fitness in *E. coli* ST131 using phylodynamics

**DOI:** 10.1101/2024.06.10.598183

**Authors:** Lenora M. Kepler, Manuel Jara, Bejan Mahmud, Gautam Dantas, Erik R. Dubberke, Cristina Lanzas, David A. Rasmussen

## Abstract

Antimicrobial resistant pathogens such as *Escherichia coli* sequence type 131 (ST131) pose a serious threat to public health globally. In the United States, ST131 acquired multiple antimicrobial resistance (AMR) genes and rapidly grew to its current high prevalence in healthcare settings. Notably, this coincided with the introduction and widespread use of antibiotics such as fluoroquinolones, suggesting AMR as the major driver of ST131’s expansion. Yet, within ST131, there remains considerable diversity between strains in resistance profiles and their repertoires of virulence factors, stress factors, plasmids, and other accessory elements. Understanding which genomic features contribute to ST131’s competitive advantage and their relative effects on population-level fitness therefore poses a considerable challenge. Here we use phylodynamic birth-death models to estimate the relative fitness of different ST131 lineages from bacterial phylogenies. By extending these phylodynamic methods to allow multiple genomic features to shape bacterial fitness, we further quantify the relative contribution of individual AMR genes to ST131’s fitness. Our analysis indicates that while many genomic elements, including various AMR genes, virulence factors, and plasmids, have all contributed substantially to ST131’s rapid growth, major increases in ST131’s fitness are largely attributable to mutations in gyrase A that confer resistance to fluoroquinolones.

**Author summary:** ST131 is a pandemic lineage of *E. coli* that has spread globally and is now responsible for a large percentage of blood and urinary tract infections that cannot be treated with many common antibiotics. While antibiotic resistance has undoubtedly given ST131 a competitive edge, the relative importance of resistance compared with other factors shaping a pathogen’s growth or transmission potential (i.e. fitness) is often difficult to measure in natural settings. Here, we present a method that allows us to look at the entire spectrum of factors determining a pathogen’s fitness and estimate the individual contribution of each component to pathogen’s overall fitness. Our results suggest that resistance to fluoroquinolones, a widely used class of antibiotics, provides ST131 with a disproportionately large fitness advantage relative to many other factors with more moderate fitness effects. Understanding what determines the fitness of ST131 therefore provides insights that can be used to curb the spread of resistance and monitor for emerging lineages with high pandemic potential due to shared fitness enhancing attributes.

## Introduction

The US Centers for Disease Control and Prevention (CDC) has named antimicrobial resistance (AMR) as “one of the greatest global public health challenges of our time” [1]. Infections with resistant bacteria are associated with treatment failure, high mortality, prolonged hospital stays, and high healthcare costs [1]. These infections are commonly caused by organisms like *Escherichia coli* (*E. coli*) that have acquired the ability to produce extended-spectrum beta-lactamase (ESBL) enzymes and thus cannot be treated with commonly used beta-lactam antibiotics such as cephalosporins. In 2017, ESBL-producing Enterobacteriaceae (ESBL-E) caused an estimated 197,400 cases of blood or urinary tract infections and 9,100 deaths in hospital settings in the US alone [1]. ESBL-E can also cause disease in healthy individuals without prior healthcare exposure, and based on recent surveillance studies, the total burden in both healthcare settings and community is likely to be much higher than current estimates [2, 3].

*Escherichia coli* (*E. coli*) sequence type 131 (ST131) is the most frequently isolated ESBL-E in blood and urine infections [2] and has played a major role in the rapid increase and global dissemination of ESBL-Es [4, 5]. A single clade of ST131, clade C, has driven the explosive growth of the sequence type [6]. Clade C is distinguished by high-level chromosomal fluoroquinolone resistance acquired shortly after fluoroquinolones came into wide clinical use [7, 8]. Two of clade C subclades have also acquired and stably maintained ESBL genes: clade C2 has long carried *bla*CTX-M-15 while clade C1 more recently acquired *bla*CTX-M-27 [5], spawning the birth and global spread of sub-clade C1-M27. In contrast, the more ancestral clade B and the more divergent clade A have been broadly susceptible to both fluoroquinolones and beta-lactams and have historically accounted for fewer sampled isolates. However, several recent studies have described rapid growth of clade A subclones after acquiring both chromosomal fluoroquinolone resistance and the CTX-M-27 ESBL gene.

While these broad evolutionary patterns suggest that the competitive advantage of different ST131 clones is driven at least in part by the acquisition of key resistance genes, the extent to which AMR is responsible for ST131’s fitness advantage remains unknown [9]. Indeed, while these AMR elements undoubtedly confer major fitness advantages in clinical settings where antibiotics are extensively used, they may come at a cost in settings with lower antibiotic pressure [10–12]. Of note, unlike some other common healthcare-associated pathogens, ST131 is frequently transmitted in community settings and can cause infections in otherwise healthy individuals [13, 14]. In addition, gut colonization can precede infection and populations often evolve for months or years in the intestinal tract before colonizing extra-intestinally [15–17]. Thus, the success of ST131 may also in part be due to virulence and stress factors that facilitate its colonization and persistence in the gut niche.

While much work has been done to elucidate the factors driving ST131’s competitive advantage, these studies often focus on the effects of a single a genomic feature under controlled environmental conditions and within the context of an isolated bacterial colony [18–21]. In contrast, phylodynamic methods offer a means to explore the genomic determinants of fitness in terms of a pathogen’s growth rate at the population-level [22]. Because pathogen phylogenies are strongly shaped by the evolutionary and epidemiological processes that generate them, it is possible to learn about these processes from the topology or branching structure of phylogenetic trees [23]. For example, more fit pathogen lineages that transmit at a higher rate will, on average, branch more often and give rise to more sampled descendants [24]. Previous phylodynamic work has therefore been able to quantify the relative fitness of antibiotic resistant versus sensitive strains of bacteria [25–27]. However, these studies have primarily focused on the impact of a single pathogen trait, whereas in reality bacterial fitness likely reflects a complex interplay of numerous factors, including diverse genomic elements, environmental pressures, and host-pathogen interactions. In previous work [28], we therefore extended birth-death phylodynamic models to allow many factors to shape pathogen fitness by using a function that maps the presence or absence of many fitness-related features to a lineage-specific transmission rate. With this approach, we can estimate the relative fitness of different lineages while also determining the relative importance of each feature to total fitness. Moreover, by allowing many features to impact fitness, we can account for potentially confounding background effects that may otherwise cause fitness effects to be erroneously attributed to the wrong feature.

Here, we apply our phylodynamic framework to learn how multiple antibiotic resistance genes as well as other genomic features jointly shape the fitness of *E. coli* ST131. Using clinical isolates obtained from human blood and urine samples collected in the United States, we first estimate the relative overall fitness of distinct ST131 lineages circulating within this population. We then decompose the fitness into individual fitness components to quantify the relative contribution of AMR genes and other genomic features to overall fitness. Subsequently, we further tease apart the contribution of individual AMR genes, plasmids, and virulence and stress factors, allowing us to quantify their individual contributions to overall fitness. Collectively, these analyses allow us to determine the most important determinants of ST131’s fitness in its natural transmission environment.

## Results

### Phylogenetic history of ST131

In order to interrogate the dynamics of ST131 fitness, we examined publicly-available blood and urine samples that were taken in the United States from humans in clinical settings. After quality control (see Methods and S1 Table), this dataset encompassed 883 samples from across the United States with collection dates spanning from to 1985 to 2022 (see S1 File). We first reconstructed the evolutionary history of our sampled strains as a time-calibrated phylogeny from a core genome single-nucleotide polymorphism (SNP) alignment (Fig 1). The maximum-likelihood tree inferred from our dataset is consistent with the previously described evolutionary history of ST131, and our estimates of clade divergence times are also consistent with prior Bayesian analyses of international isolates [7, 16]. The most recent common ancestor (MRCA) of all samples is estimated to have arisen in 1862, when clades A and B diverged. We estimate that clade B0 diverged from clade B in 1963; C0 from B0 in 1966; C1 from C0 in 1981; C2 from C1 in 1991; and C1-M27 in 2001 (Fig 1).

**Fig 1.**
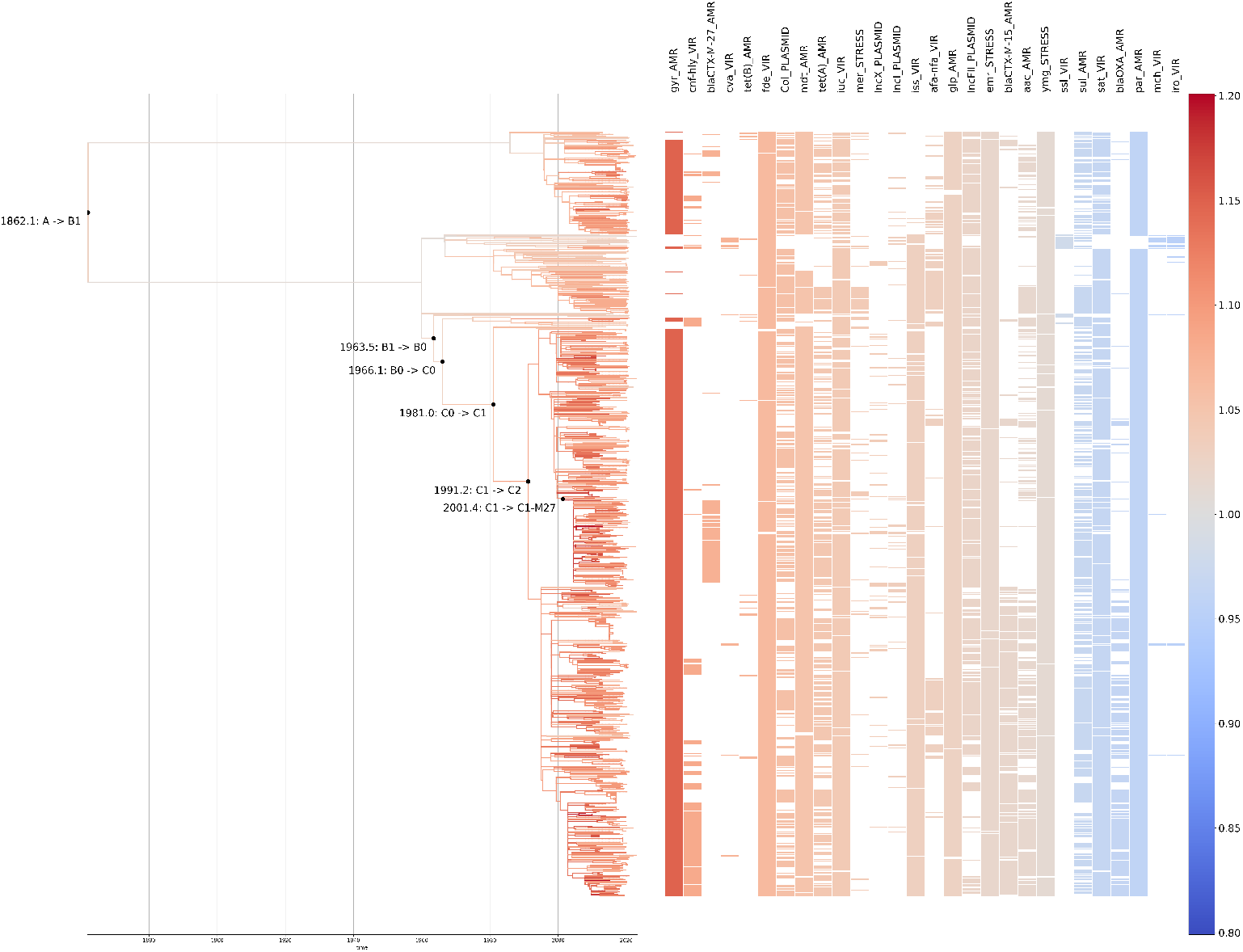
Maximum-likelihood time-calibrated ST131 phylogeny. Branches are colored by their estimated fitness. Distribution of significant features across samples in the phylogeny is displayed on the right. Colored bands, representing the presence of a feature in a sample, are also colored according to their estimated fitness impact.

While the overall clade distribution is consistent with previous work noting the dominance of clade C and especially C2 [6], we note an increasing percentage of isolates from clade A beginning around 2010 (Fig 2), accounting for more than 25% of samples by 2020. Other recent longitudinal sampling efforts have shown a similar pattern of increasing growth within Clade A [29, 30].

**Fig 2.**
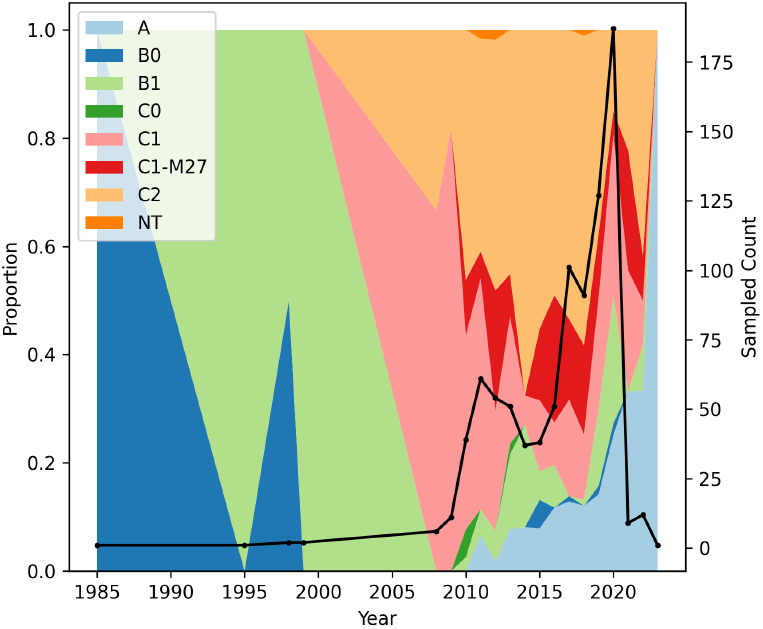
Proportion of sampled ST131 isolates from each clade by year. Black line indicates total number of samples per year.

### Overall ST131 fitness through time

We model the fitness of ST131 as a combination of the effects of genetic features (AMR genes and AMR-conferring point mutations, plasmids, virulence factors, and stress-related genes) while also taking into account the confounding effects of variable sampling effort and background transmission rates through time (see Methods). Here, fitness is assumed to be proportional to a lineage’s growth rate in the host population. We can thus estimate the overall relative fitness of each lineage in the phylogeny as the product of the effect of each of the genetic features present in the lineage.

Averaging over all lineages in the phylogeny present at a given interval, there has been an overall increase in ST131’s fitness over time (Fig 3). This fitness increase appears to be largely driven by the acquisition of AMR. Grouped together, we infer that on average, the AMR elements that early lineages carried (*parC/E, sul*, Fig 5), actually had a net negative effect on fitness. However, starting around 1980, lineages began accumulating beneficial AMR mutations. Collectively, AMR features are responsible for much of the increase in ST131’s overall fitness and the average fitness advantage they confer has risen steadily over time (Fig 3). In contrast, the average fitness contributions of virulence and stress factors appear more constant over time. These results strongly suggest that AMR features have collectively increased the average growth rate of ST131 through time.

**Fig 3.**
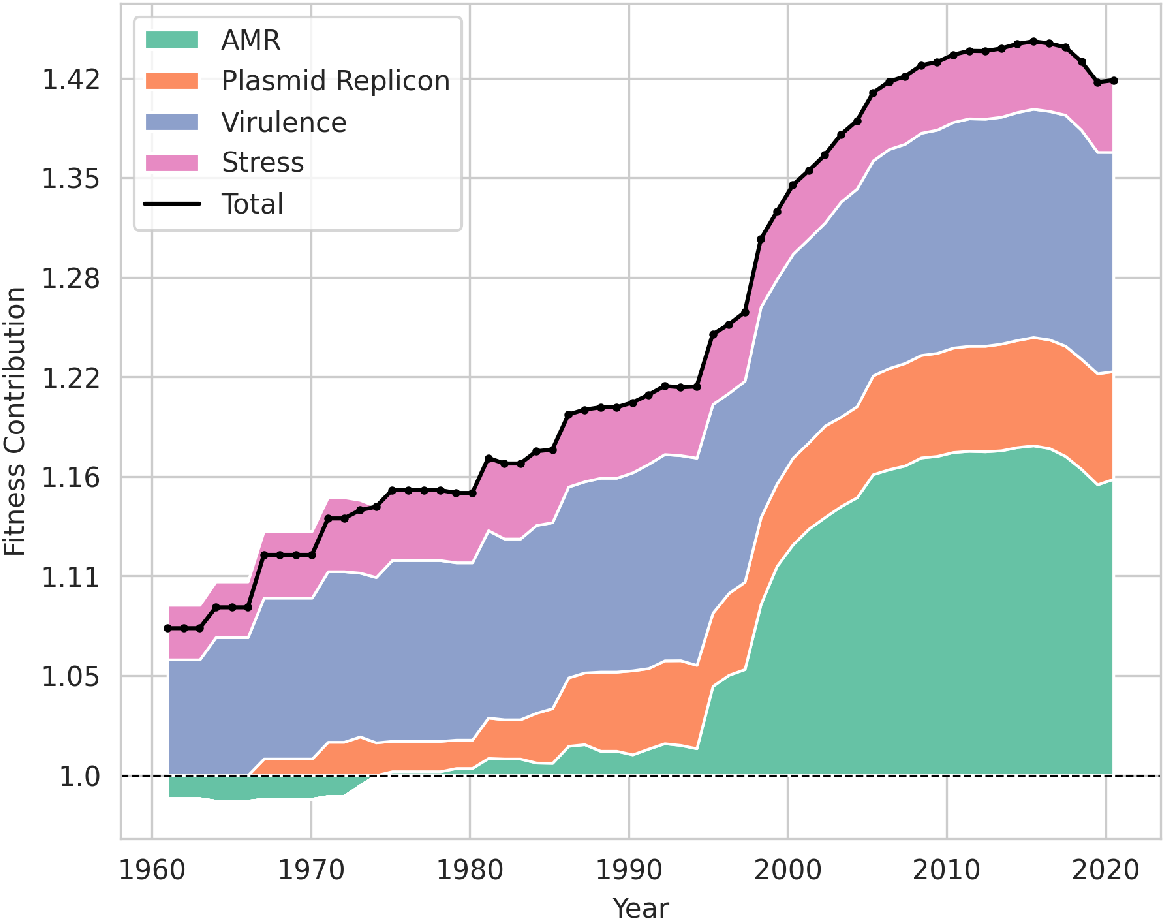
Changes in total ST131 fitness due to different components of fitness through time. The black line indicates total fitness averaged over all lineages present in the phylogeny at each time point. Colored regions indicate the estimated contribution of each fitness component to total fitness over time. Those that are below one indicate negative impacts. The contribution of each fitness component was determined by first computing for each lineage the summed fitness effect of all features belonging to each fitness component and then averaging these component-wise effects across all lineages present at each time point.

Consistent with the large fitness benefit attributable to AMR, the historically more drug-resistant clades C2 and C1-M27 [4, 6, 31, 32] are overall the most fit, with a 1.17x and 1.19x fitness advantage over clade B1, respectively. The relative fitness of clade A compared to B1 is 1.11x, while C1 clades without CTX-M7 are only 1.03x as fit as B1.

### Estimated fitness effects of genetic features

Many potentially fitness-effecting genetic elements have been independently acquired or lost across multiple different lineages in the ST131 phylogeny (Fig 1). The presence of these features across different genetic backgrounds allows us to infer the contribution of individual AMR, plasmid, and virulence features separately from the overall fitness of the lineages in which they occur. We assume that the presence of a given gene or mutation scales the overall fitness such that a fitness effect greater than one is beneficial and those less than one are deleterious. After imposing L1 regularization to avoid overfitting, 26 of 67 genetic features (about 39%) were estimated to have no impact on fitness. Of the remaining 41 features, 13 had 95% confidence intervals overlapping one and were thus determined not to have a significant effect (Table 1). Confidence intervals varied greatly, with the most uncertainty occurring for features that are less prevalent. In total, we estimate that 28 (about 42%) of the genetic features analyzed have a significant impact on the fitness of ST131 lineages in our dataset. These impacts are mostly beneficial, with only seven (25% of all significant effects, 10% of total) decreasing fitness (Fig 4).

**Table 1.**
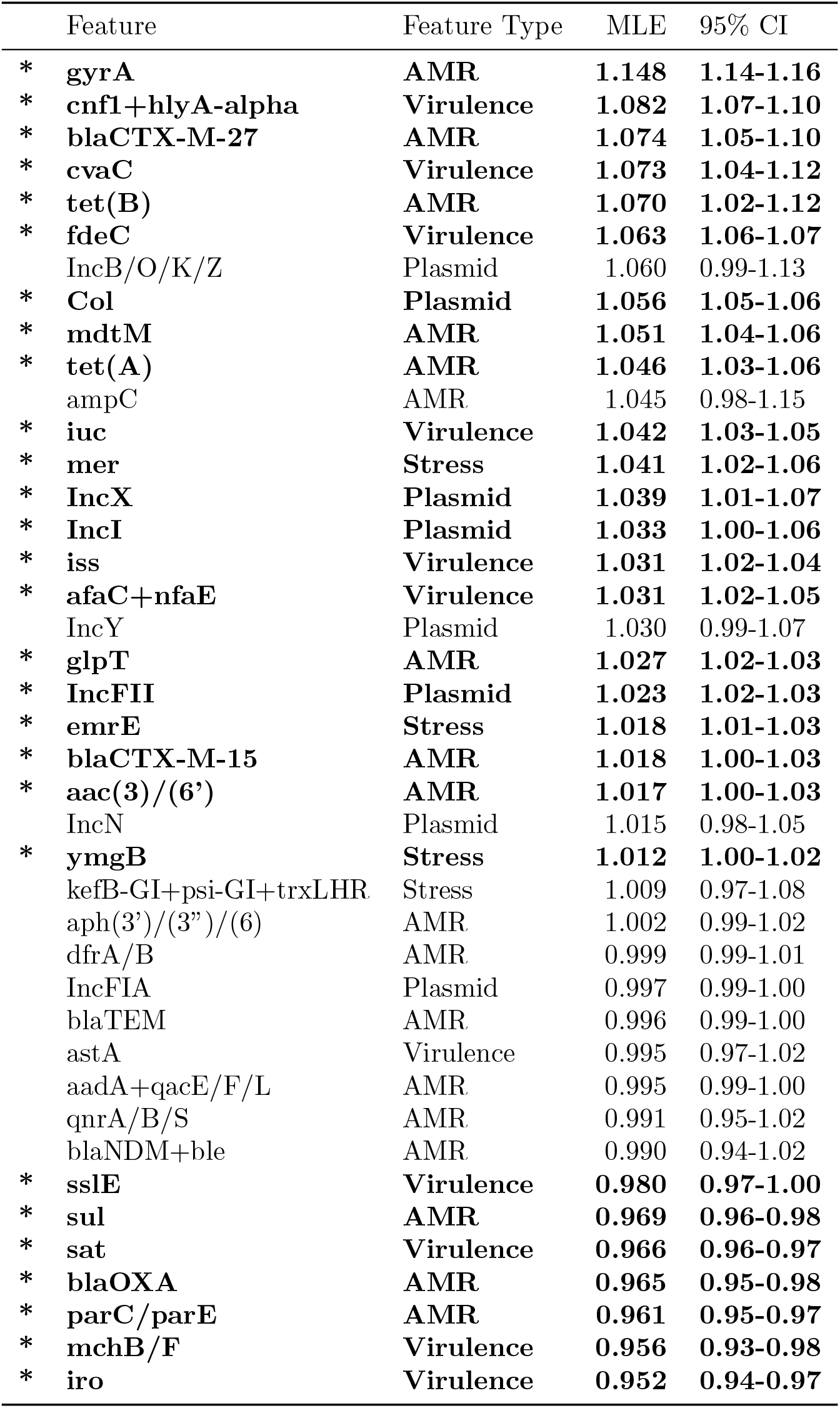
Maximum-likelihood estimates of genetic features that were not dropped out of the model. Significant features are bolded with an asterisk.

**Fig 4.**
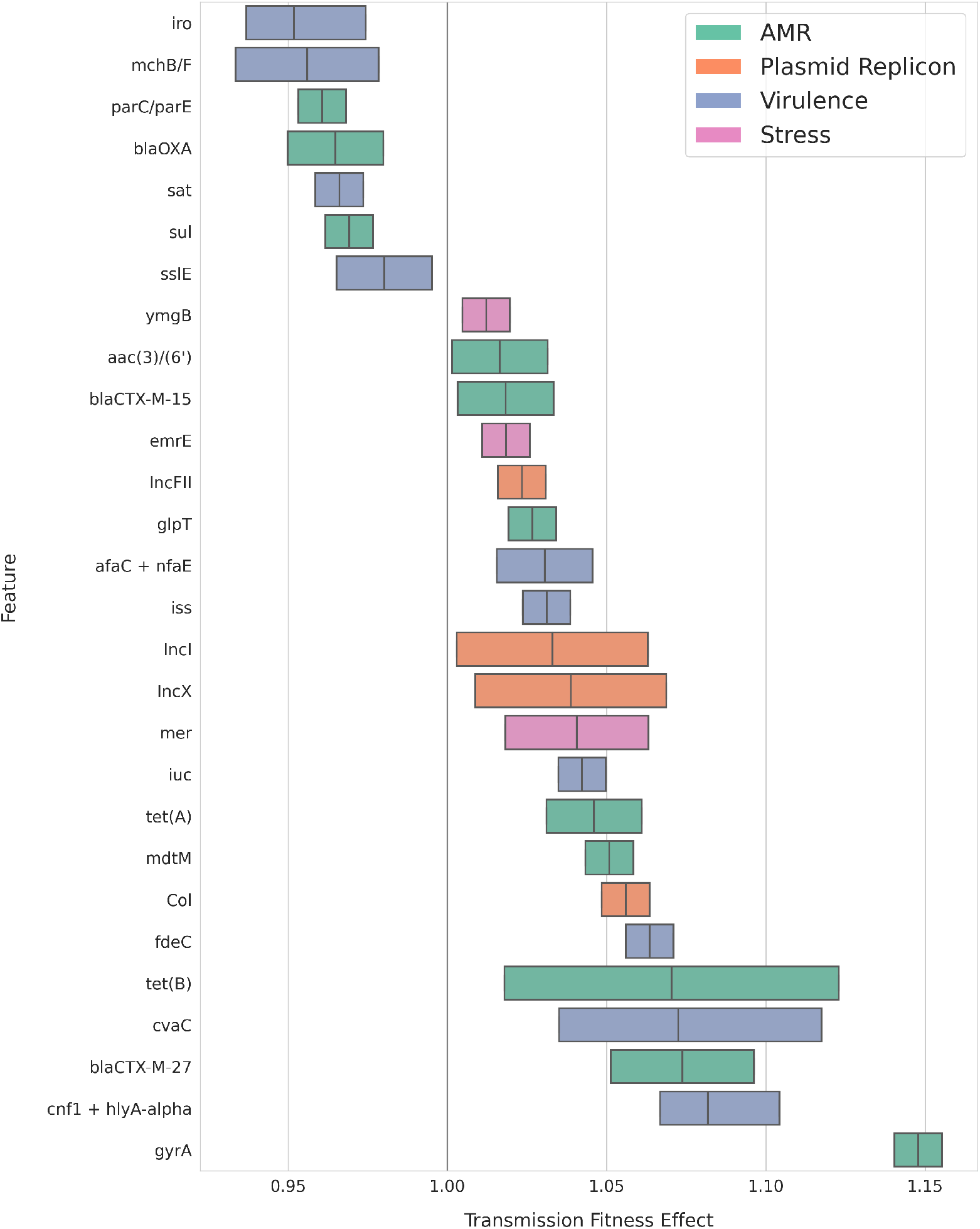
Genetic features with significant estimated fitness effects. Bars cover the 95% confidence interval are colored by feature category. The middle line represents the maximum likelihood estimate. Colors show how features are grouped into different categories.

By far, the largest fitness advantage to ST131 comes from the acquisition of one of two amino acid substitutions on the chromosomal gyrase A (*gyrA*) gene, (S83L/*gyrA-1A* or D87N/*gyrA-1B*). The presence of one or both of these fluoroquinolone resistance mutations increases a lineage’s base fitness by a factor of 1.15x. Indeed, the acquisition of these mutations by a C-clade ancestor in the early 1980s appears to be the greatest source of increased fitness in clades C1 and C2 (Fig 5). Subsequently, these *gyrA* mutations have been independently acquired by lineages in clade A (around 1996) and B1 (1997 to 2003) (S2 Fig), and are now the largest contributor of fitness to clade A.

**Fig 5.**
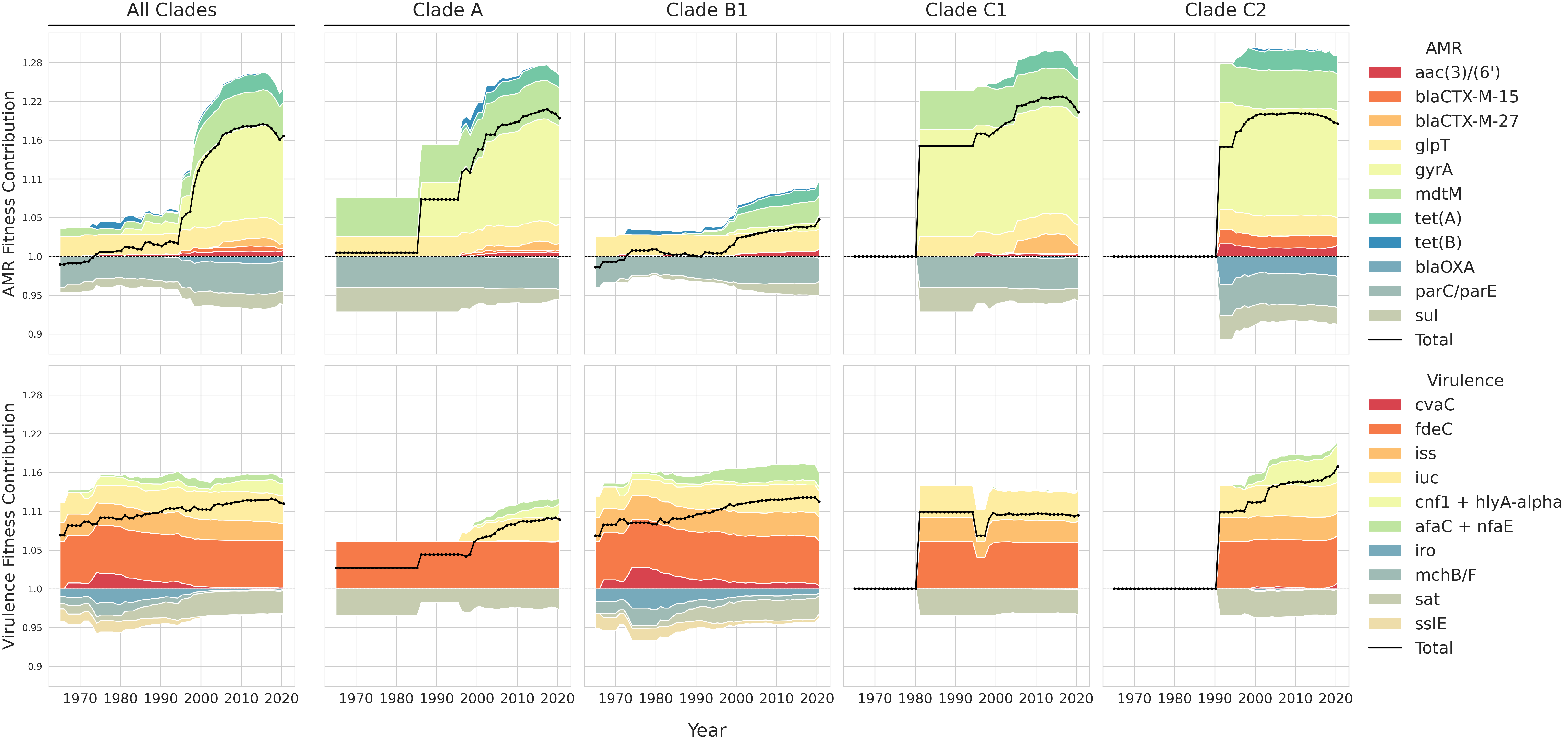
Fitness contributions of AMR and virulence through time. Comparison of AMR (top) and virulence (bottom) contributions to the overall fitness of ST131 as a whole (leftmost column) and to the fitness of major ST131 clades over time. At each time interval, the amount of fitness attributable to each feature is summed across all extant lineages in a given clade and then averaged, giving a measure of a feature’s fitness effect weighted by its prevalence within a clade.

Other AMR features with positive estimated fitness effects appear to have similarly driven fitness increases. In fact, the increase in ST131’s fitness starting in the mid-1990s appears to be driven almost entirely by the acquisition of AMR-conferring elements. In addition to *gyrA*, multidrug resistance protein *mdtM* E448K (MLE fitness effect: 1.05) and tetracycline resistance gene *tet(A)* (MLE fitness effect: 1.04) were convergently acquired across multiple clades and substantially increased overall ST131 fitness (Fig 5). To a lesser and more clade-variable extent, ESBL genes *bla*-CTX-M-27 (MLE fitness effect: 1.07) and *bla*-CTX-M-15 (MLE fitness effect: 1.02) as well as the aminoglycoside gene aac(3)/(6’) (MLE fitness effect: 1.02) also contributed to ST131’s fitness.

In contrast to AMR genes that contribute to variation in fitness across ST131, the fosfomycin resistance mutation *glpT* E448K [33] was present ancestrally and has been lost only a few times. Thus, despite a relatively small fitness benefit of 1.027, it contributes substantially to the sequence type’s overall fitness.

Despite the comparatively small and stable contribution of virulence factors towards overall fitness through time (Fig 5), the second largest fitness effect comes from toxin-encoding virulence factors *cnf1* (Cytotoxic Necrotizing Factor 1) and α*-hly* (analyzed together in this dataset) that have been found to increase ST131’s ability to colonize the gut [20]. Similarly, *cvaC* (colicin V, MLE fitness effect: 1.07), a colicin that also facilitates colonization [34], and fdeC (MLE fitness effect: 1.06), an adhesin, have positive fitness effects.

We included the presence of plasmid replicon genes in our analysis to attempt to disentangle the effect of plasmids per se from the effect of the genes that plasmids carry. For instance, in many cases, the metabolic cost of plasmid carriage is deleterious, but the significant benefit of the genes it encodes makes its carriage overall beneficial [10]. However, we found this to not be the case here: no replicons had a deleterious effect on ST131 fitness. Intriguingly, Col family replicons provided the largest fitness benefit of all plasmid types analyzed (MLE fitness effect: 1.06). While often a replicon associated with small plasmids that encode colicins, in ST131 they are also found on large IncF plasmids with IncFII (MLE 1.023).

### Decomposing the components of fitness variation

Because selection acts on fitness variation at the population level [35], understanding how much fitness variation is attributable to each fitness component provides another means to quantify the relative importance of different features to ST131’s overall evolutionary dynamics. While we expect fitness to vary between lineages due to the genetic and background features included in our model, we also expect that there may be additional unmodeled sources of fitness variation. We thus extend our model to allow for the fitness of each lineage to vary due to branch-specific random effects. Doing so allows us to consider how much fitness variation is attributable to each genetic feature or other component of fitness compared to the total fitness variation from all sources.

On the whole, our modeled features are able to account for almost half (47.01%) of overall variation in fitness across the entire ST131 phylogeny. Genetic features (AMR, plasmid replicons, virulence factors, and stress factors) contribute 19.55% of variance, while time-varying and background fitness account for 27.46%. AMR features alone account for 13.68% of fitness variation across ST131 lineages.

Despite the convergent acquisition of several genetic features across multiple lineages, the overall variance in fitness has increased with time (Fig 6, left), largely due to increasing variation in branch-specific random effects (Fig 6, right). Early on, fitness differences attributable to the carriage of virulence factors and, to a lesser extent, plasmids, were responsible for much of the variation between lineages (Fig 6, right). But the importance of AMR steadily grew over time, becoming the largest differentiating factor by 1990 and increasing in importance until the early 2000s when variation in fitness from AMR plateaued and subsequent increases in fitness variation were attributable only to non-genetic features (Fig 6, left). This trend is potentially attributable to the large effect of gyrA mutations on fitness: it contributed strongly to variance when it was acquired by clade C, and then more or less plateaued when clade A acquired gyrA mutations around 1997 and most sampled features had one or both gyrA mutations (S2 Fig).

**Fig 6.**
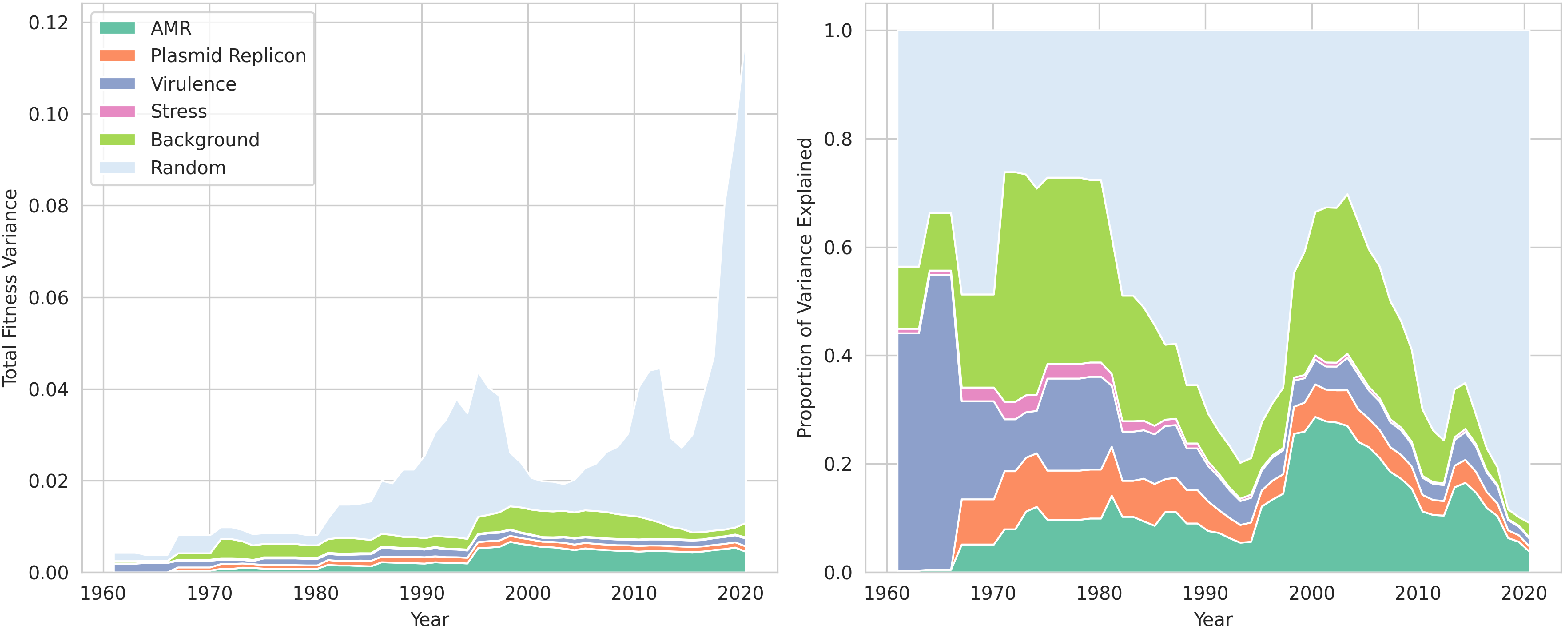
Contribution of each model component to fitness variation over time. Left: Amount of variance accounted for by each category over time. Right: Proportion of total variance explained by each category over time.

**Fig 7.**
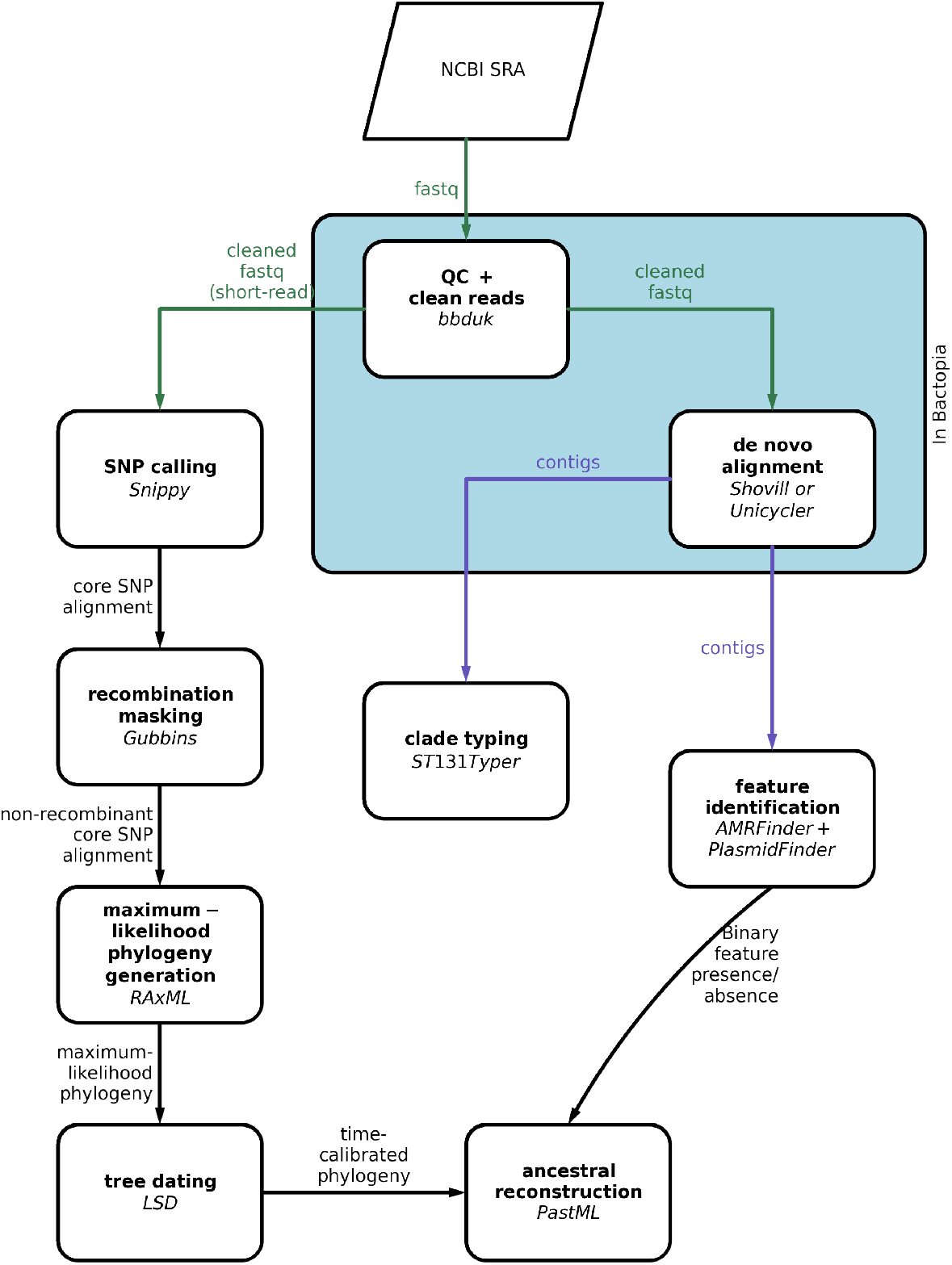
Workflow for phylogenetic and ancestral feature reconstruction. Steps inside blue box were performed using the Bactopia workflow.

## Discussion

Understanding what factors shape pathogen fitness presents a daunting yet critical challenge as understanding what shapes fitness in real-world settings is tantamount to understanding what factors drive the growth and epidemic potential of different lineages. For example, while AMR has long been suspected to a be a key determinant of ST131’s fitness [6, 9], the fitness effects of individual AMR genes relative to other components of bacterial fitness has been difficult to quantify. Our analysis of ST131 fitness highlights the benefits of phylodynamic methods that allow the fitness effects of multiple features to be quantified and compared at the population-level. In particular, we were able to disentangle the relative contribution of many genetic features including AMR elements, virulence factors, and plasmids to overall fitness, while controlling for other background sources of fitness variation, enabling us to gain a more holistic understanding of what factors drive the spread of ST131 in its natural setting.

Overall, the genomic features we analyzed can explain a striking amount of fitness variation within ST131. Specifically, genes and mutations that confer antimicrobial resistance seem to have been the dominant force shaping the fitness and evolutionary trajectory of ST131 and, until the early 2000s, the amount of variation attributable to these resistance features has increased over time. Substantial fitness gains have come from fluoroquinolone resistance mutations, ESBL elements, and tetracycline resistance genes. In addition to these AMR features, acquisition of Col plasmids and colonization-promoting virulence factors like *cnf* /α*-hly, cvaC*, and *fdeC* strengthened the competitive advantage of successful ST131 lineages.

Our analysis indicates that the presence of chromosomal mutations to the *gyrA* gene are by far the most important determinant of ST131 fitness. While these mutations (D87N and S83L) have been recognized as both clinically and evolutionarily important to ST131 [7–9], the magnitude and persistence of their fitness impact is nonetheless surprising. While the emergence of ST131 as a pandemic lineage occurred at the same time that fluoroquinolones came into widespread use [7], recent years have seen a plateau (2011 through 2015) and then sharp decline in their prescription [36] following both a push to curb AMR by decreasing their prescription for uncomplicated urinary tract infections [37] and the addition of a black-box warning about their potential side effects [38]. Given that the *gyrA* mutations have not decreased in frequency since fluoroquinolone use declined, this suggests that *gyrA* mutations may be beneficial even without the widespread use of fluoroquinolones in clinical settings.

Indeed, numerous studies have shown that across bacterial species, *gyrA* mutations can be beneficial even in the absence of antibiotics [29, 39–41]. However, the reasons behind this benefit remain unclear. Some evidence supports the idea that mutations in gyrA alter the supercoiling of DNA, leading to potentially adaptive changes in gene expression during the stress response [39, 42, 43]. *gyrA* mutations may also allow for more stable, lower-cost carriage of IncF plasmids by increasing the efficiency of plasmid replication [9, 44, 45], though a recent study failed to see this effect in vivo [46]. Yet another line of evidence suggests that a non-serine residue at position 83 in *gyrA* may be overall more evolutionarily favorable across species, but that ancestral bacteria evolved to express that amino acid because it confers resistance to the natural antibiotics that would have been present in their ancestral environments [47, 48]. Thus, rather than an evolutionary change that allows *E. coli* to persist in fluoroquinolone-rich environments, *gyrA* S83L may be better thought of as the reversion of a metabolically costly mutation that is no longer necessary for bacterial survival (see discussion in [49] and [50]).

The recent surge of *E. coli* sequence type ST1193, which also shares these key mutations, validates their importance to population-level fitness [51, 52]. In fact, ST1193, first isolated in only 2011 [53, 54], appears to be the only *E. coli* lineage that is able to out-compete ST131. In a large US study, ST1193 was responsible for 31% of *E. coli* infections that occurred from October 2019 to September 2020, second only to ST131 [55]. Altogether, these results suggest that antibiotic stewardship alone may not be enough to curb the spread of fluoroquinolone resistance among *E. coli*, as some of the mechanisms of resistance are adaptive even absent the antibiotic.

Several other AMR genes were found to be beneficial to ST131, including ESBL genes *bla*CTX-M-15 and *bla*CTX-M-27. However, given the sharp rise in ESBL-E prevalence and the global spread of ESBL-associated ST131 clades C2 and C1-M27, the relatively small magnitude of their effect sizes (1.07x for M-27 and 1.02x for M-15, Fig 4) is surprising.

Given the immense fitness benefit conferred from AMR genes, it could be theorized that the stable carriage of even large, F-type plasmids characteristic of ST131 is entirely due to the carriage of these genes offsetting an intrinsic fitness cost of carrying these plasmids. However, we find that, independent of AMR and virulence content, there is no evidence that any plasmid type has an inherent fitness cost to ST131. In fact, several plasmid types (Col, IncX, IncI, and IncFII) are estimated to be independently beneficial. These population-level results validate previous experimental work demonstrating that there are no costs associated with plasmid carriage in ST131 [19, 56], or that these effects only occur under specific environmental and metabolic circumstances [57].

Recent work has found that IncF-family plasmids are enriched in conjugation and plasmid maintenance genes (e.g. toxin/antitoxin pairs), which may facilitate the maintenance of these plasmids and may even help drive the expression of ESBL genes, both of which could positively impact ST131 fitness [58].

While the majority of ST131’s fitness seems to be driven by AMR, virulence factors have also played a major role in shaping ST131’s evolution. Virulence factors that enhance fitness seem to be those that allow ST131 to colonize and persist, like *cnf* /α*-hly, cvaC*, and *fdeC*, rather than those that increase disease severity. These results are consistent with recent hypotheses that the main environment in which ST131 thrives, evolves, and transmits is the gut of individuals in the community, not necessarily in antibiotic-rich hospital settings [9, 20, 59, 60]. Pitout and Finn [61] further suggest that this unique virulence profile may allow these lineages to persist in, and move between, different hosts and environments where they have more opportunity to be exposed to, and subsequently acquire, a diverse range of plasmids and other mobile genetic elements that allow them to rapidly adapt to new environments and outcompete the resident microbial population [62]. In fact, [63] suggest that virulence factors that enhance infection severity have been negatively selected against in clade C, even at the cost of their ability to outcompete other lineages. However, this reduced competition ability in vivo may actually lead to enhanced population level fitness. We recapitulate those findings here, where the *iro* operon, *mchB/F*, and *sslE* all significantly contribute to the reduced population-level fitness of clade B.

In addition to fitness, pathogen phylogenies can be strongly shaped by sampling biases [64–66]. The major risk here is that oversampling of certain genotypes or lineages could produce a higher branching rate that would be difficult to distinguish from higher pathogen fitness [67–69]. This is a particular concern as our ST131 samples largely came from studies (here grouped by NCBI Bioprojects) that were specifically looking for either fluoroquinolone or beta-lactam resistance. We therefore tried to account for sampling biases by including Bioproject-specific fitness effects in our model so that oversampling of specific lineages by individual studies would not bias our estimates of the fitness effects of individual features. Nevertheless, we may have overestimated the fitness effects of features that were systematically oversampled across studies. However, if there was a systematic bias, we would expect to see large fitness benefits for all overrepresented AMR features, whereas we did not infer large fitness benefits for ESBL and other AMR genes as might have been expected if there was such a strong systematic bias. We also note that our analysis focused only on clinical isolates of ST131, and so our results may not generalize to lineages circulating outside of clinical environments, where different genetic features may play a role in determining pathogen fitness. More representative sampling of both susceptible and resistant genotypes in and outside of healthcare settings would therefore provide a more complete picture of the factors shaping ST131’s long-term fitness.

While our results support the importance of AMR in the success of ST131, they raise other questions about the nature of AMR’s benefit. Rather than bolstering the fitness of a lineage purely by conferring antibiotic resistance, the benefit of these genes may not be entirely dependent on the presence of antibiotics [70]. This will be an important distinction to untangle moving forward. While increased antibiotic stewardship and vigilance in clinical settings would significantly curb the spread of multi-drug resistant pathogens like ST131 if AMR was largely deleterious outside of an antibiotic-rich context, curbing its benefits in the context of often non-symptomatic individuals in the diffuse community setting may require very different tactics. Using our phylodynamic framework, specific hypotheses about what additional factors promote or constrain the growth of AMR in and outside of antibiotic-rich settings settings can be tested.

## Methods

### General approach

We aim to estimate the relative fitness of any ST131 lineage from the branching structure of the pathogen phylogeny. Here, fitness is defined in terms of a lineage’s exponential or intrinsic growth rate *r* = λ_*n*_ − μ_*n*_, where λ_*n*_ and μ_*n*_ are the lineage-specific birth and death rates, respectively. Given a set of binary features 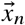 used to predict fitness, we parameterize the birth rate λ_*n*_ as a multiplicative fitness model where each feature *i* scales a lineage’s fitness by a factor of *β*_*i*_ when present, while accounting for time-varying non-genetic effects with a series of time-interval-specific effects λ_*T*_ :

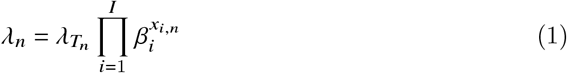

Evaluated on a log scale, this fitness function resembles a traditional linear regression problem. However, because we do not directly observe a lineage’s fitness, we cannot find a best-fitting approximation in the usual fashion, e.g. by minimizing least squares.

Instead, we take advantage of the branching structure of a phylogeny 𝒯 generated under a birth-death-sampling process [71, 72]. The likelihood that the phylogeny evolved as observed under this model is determined by the rates at which lineages are born via transmission, die or are removed from the infectious population, and are sampled. We can therefore find the lineage birth rates that are most likely to have generated the observed topology. That is, we find the values of each coefficient *β*_*i*_ and time effect λ_*T*_ that maximize the likelihood of obtaining the observed phylogeny given the fitness function in (1). Assuming there is a constant death rate shared among all lineages, the estimated birth rates will be proportional to a lineages relative fitness or growth rate *r*.

### Data acquisition and availability

We downloaded all *E. coli* assemblies from NCBI’s Isolates Browser that were from human clinical urine or blood specimens (N=13,017). To facilitate expedient in-silico sequence typing, we chose to first download pre-assembled genomes. After removing duplicate BioSamples, we used the software package MLST [73] to identify each isolate’s sequence type using the PubMLST Achtman sequence definitions [74]. We extracted all isolate sequences classified as ST131 (N=3,659) and further narrowed the dataset down to those ST131 isolates from the USA that had a valid collection date, raw reads available, and for which there were at least five samples in the dataset for its BioProject (N=900).

Once these isolates were identified, we then downloaded the associated raw read files from the NCBI Sequence Read Archive (SRA) using SRA Read Downloader [75]. If a BioSample had both long and short-read sequencing, both reads were downloaded in order to perform hybrid assembly. In the case where short-read sequencing was performed multiple times, we downloaded only the most recent set of reads.

The scripts used to perform analysis, are available on GitHub (https://github.com/lenorakepler/ST131Fitness. Accession numbers and metadata for our dataset are also available in the repository, as well as in S1 File.

### Phylogeny and feature reconstruction

#### Assembly and phylogenetic reconstruction

Quality control and genome assembly were performed using the Bactopia version 2.2.0 pipeline [76]. The pipeline performs quality control, adapter trimming, and error correction on raw Illumina reads using various software including FastQC [77], BBTools [78], and Lighter [79]. High-quality, cleaned reads are then assembled de novo. Isolates with only short reads were assembled using Shovill [80] whereas isolates with both short and long reads were assembled with Unicycler [81].

To find the set of shared, clonally inherited genomic sites with which to build the phylogeny, cleaned short reads were aligned against the high-quality ST131 reference EC958, a 2014 urine isolate from the United Kingdom that belongs to clade C2 [82] using Snippy [83]. Freebayes within Snippy was then used to call variants and output the core genome (here, 97,345 genomic sites that are present in all samples, of which 15,655 are polymorphic). Probable sites of recombination were masked using Gubbins [84], leaving 7,445 clonally-inherited core SNPs [85].

RAxML [86] was used to infer a maximum likelihood tree using a GTRCAT approximation of rate heterogeneity. LSD version 0.3 beta in constrained mode [87] was then used to estimate the root position and date the phylogeny.We subsequently fixed the reconstructed phylogenetic tree topology and branching times for all downstream analyses.

We identified the presence of AMR genes and mutations, virulence factors, and stress-related genes on our assembled contigs with NCBI’s AMRFinder+ [88] with organism “Escherichia” and including the “plus” proteins. The presence of plasmid replicon types were similarly identified from our contigs using staramr [89], which blasts against the PlasmidFinder database, [90] using default parameters with organism “escherichia coli”. Isolate ST131 clade types were inferred using ST131Typer (https://github.com/JohnsonSingerLab/ST131Typer), an in silico implementation of the allelic profiling and categorization developed in [91].

Features were grouped to the level of gene, gene cluster, or plasmid replicon type and coded as present in an isolate if any of the genes in the group were found. We further subdivide IncF replicon types because of their overall abundance in human *E. coli* isolates [92]. After performing this feature grouping, the covariance between features was analyzed (see S1 Fig) and features with greater than 0.95 covariance were considered as one feature.

We also included in our model several features intended to account for the confounding effects of varying sampling effort across and time and between studies. We created three time intervals, estimating a separate background effect on fitness for each interval to account for increased sampling over time. We also included features to account for the background effects of different bioprojects (one-hot-encoded, to account for differing sampling intensity and environmental contexts across isolates) and specimen types (urine, 1, or blood, 0), to prevent differential collection or gene content from biasing our estimates of genetic fitness.

After extracting feature information about our sampled isolates, we use the known tree topology to reconstruct the probable feature states of unsampled (ancestral) nodes on the tree. Features were converted into binary format with 1 representing the presence of the feature and 0 denoting its absence. We then reconstructed ancestral feature states using PastML [93].

### Phylodynamic inference

#### The phylodynamic birth-death-sampling model

We divide the phylogeny along each edge *ε*_*n*_ into lineage segments *τ*_*n*_ with features 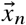 and birth-death-sampling parameters *θ*_*n*_ starts at time *t*_*s*_ and ends at time *t*_*e*_ with an event (node) that shares the same parameters. The event can represent either a feature or time-interval change (single descendant), a transmission event (two descendants), or sampling upon removal (a terminal node). The likelihood of each tree segment *τ*_*n*_ evolving as observed is the likelihood of its edge evolving as observed multiplied by the likelihood of its associated event:

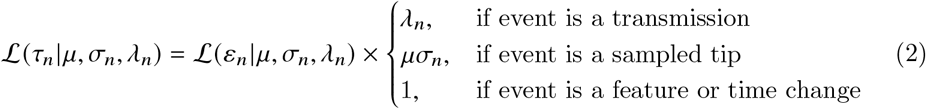

The likelihood of each edge evolving as observed ℒ (*ε*_*n*_|μ, *σ*_*n*_, λ_*n*_) given the death rate μ, the edge’s sampling fraction *σ*_*n*_ and the edge’s birth rate λ_*n*_ can be computed analytically as described in [71]. Please note that for simplicity’s sake, we describe the likelihood of edge *ε*_*n*_ as being dependent only on a single sampling probability *σ*_*n*_ and a single birth rate λ_*n*_, but because the likelihood of *ε*_*n*_ takes into account the probability that this edge produced no sample descendants, it also takes into account the sampling probabilities and any time-dependent fitness effects that would act upon descendants occurring from the edge’s event time to the present.

The probability of the entire phylogeny evolving as observed is then the product of the likelihood of each segment:

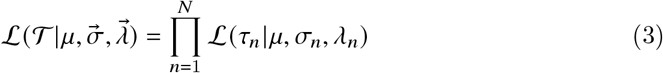

#### Model parameterization

We assume that the probability of a lineage being sampled upon removal is 0.0001 while sampling for its Bioproject was ongoing (between the time of the first Bioproject sample in our dataset to the last) and 0 otherwise. This sampling fraction was chosen to reflect the relatively small number of bacterial infections that are sampled, but may be an under-or over-estimate. While this will influence absolute fitness values, it will not change the relative fitness values that we estimate. We set the removal rate to a constant value of one, which indicates an infection/persistence time of one year before resolution, treatment, or death for all lineages.

We define the absolute fitness of a lineage as its exponential growth rate *r*_*n*_ = λ_*n*_ − μ_*n*_. By making the simplifying assumption that removal rates are constant across lineages, the relative fitness of one lineage compared to another is dependent only upon the birth (transmission) rate λ_*n*_.

In turn, we parameterize a lineage’s total fitness λ_*n*_ with three components: λ_*T*_, a time-specific scalar representing the overall background (non-lineage-specific) effect of occurring at time interval 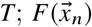, fitness due to the set of lineage-specific features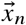 ; and *u*_*n*_, which represents the residual fitness not explainable by λ_*t*_ and 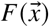. The total fitness of a lineage is then:

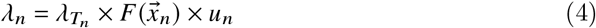

Where 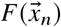 is a multiplicative fitness function where each binary feature *x*_*i*_ modifies the fitness of a lineage by *β*_*i*_ when present:

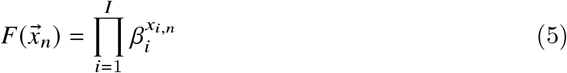

Thus, we aim to simultaneously estimate the values of both a vector of time-interval-dependant birth rates 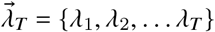 and a vector of feature-specific fitness effects 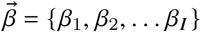.

#### Model fitting

To prevent overfitting our dataset at the expense of generality and predictive accuracy, we imposed impose an L1 regularization penalty on each the effects. Thus, we maximize the penalized log likelihood:

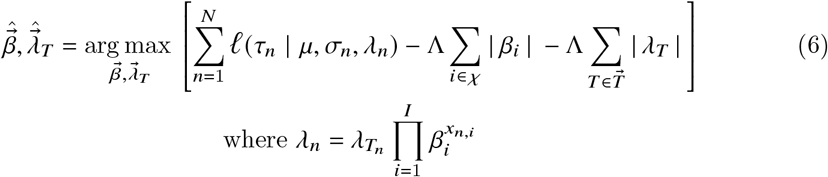

To find the optimal value of Λ, we performed four-fold cross validation by randomly dividing each edge or node of the phylogeny into four groups. For each proposed value of of Λ, we trained the model on each of the groups and then calculated the un-penalized log likelihood of the phylogeny comprised of the rest of the groups (the “test” set) given the estimated optimal 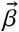 values. The value of Λ that had the highest test log likelihood across folds was chosen.

We implement our phylodynamic optimization in Python using the TensorFlow v2 library [94], which facilitates efficient mathematical computation on large-scale data. We use TensorFlow’s implementation of the Adam algorithm [95] to efficiently search for optimal parameter values by proposing new values based on their gradients.

### Statistical Analysis

#### Confidence intervals

To quantify the degree of uncertainty surrounding our estimates, we calculated 95% confidence intervals for each of the feature-specific effects *β*_*i*_. To do so, we calculated likelihood distribution by computing the penalized log likelihood for a grid of values within ∓0.75 of *β*_*i*_’s maximum likelihood estimation (MLE). The 95% confidence interval was taken to be all values of *β*_*i*_ whose distance from the MLE is less than or equal to one half of 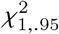.

#### Residual fitness model

To understand how much variation in fitness exists that may not be accounted for by our model, we attempted to quantify the residual fitness of each branch. To constrain these estimates, we assume a Brownian motion model of evolution. This is a stochastic process where the fitness distribution of a child branch is inherited from its parent and thus its fitness is expected to differ only based on the amount of evolution that has occurred since divergence from the parent.

To calculate random effects, we find the the residual fitness of each branch that maximizes the log-likelihood of the phylogeny, but penalize the likelihood if there is more variation in random fitness than would be expected:

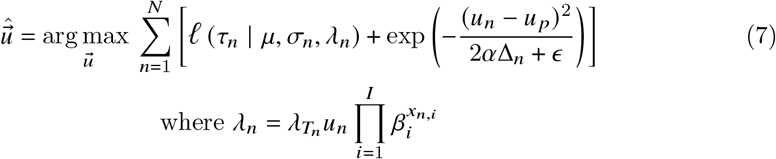

Where *u*_*n*_ is the residual fitness of branch *n, u*_*p*_ is the residual fitness of its parent, and Δ_*n*_ is the time that has elapsed since the two diverged.

We fix the effect sizes 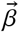 as those estimated with the previous model. The small value *∈* to prevent underflow and division by zero when α is small and parent and child lineages are very recently diverged.

The parameter α controls how much we expect a child lineage to diverge over time. We treat α as a hyperparameter that should allow enough variance between parent and child to adequately capture differences in fitness, but not so much that we are overfitting to a stochastic realization rather than the underlying distribution. To strike this balance, we find the value of α according to the hyperparameter optimization method described in [28].

#### Decomposition of fitness variance

To quantify the extent to which our genetic model accounts for variation in ST131 fitness, we partition the total variation in fitness observed through time into components attributable to our model effects and those from residual fitness not otherwise accounted for, as described in [28]. We further partition the variance explained by our model into a set of disjoint components to examine the relative impact of AMR, plasmid carriage, virulence, stress, and background factors over time.

We define the vector 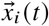 as the presence or absence of feature *i* in each of the lineages *n* alive at time *t*. Ignoring covariances among the components of fitness, the variance attributable to *c* at time *t* is simply the sum of the variances of each individual feature in *c*:

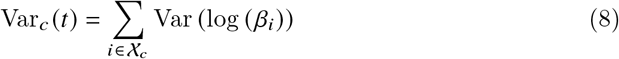

Where 𝒳_*c*_ denotes the set of all features included in fitness component *c*. Note that to ensure that the total variance attributable to *c* is not greater than the total variance, we ignore covariances. The fraction of variance attributable to component *c* at time *t* then is then simply the variance of *c* divided by the total variance, or the sum of the variance of each of the components:

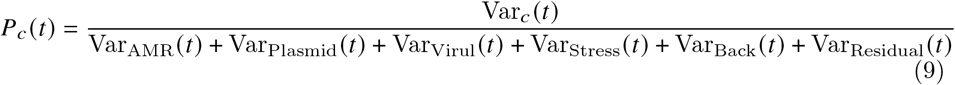

## Supporting information

### S1 Table

#### Progression of sample count

Proceeding from the top with the full data set, each row contains the criteria needed for sample inclusion along with the total number included after implementing the criteria.

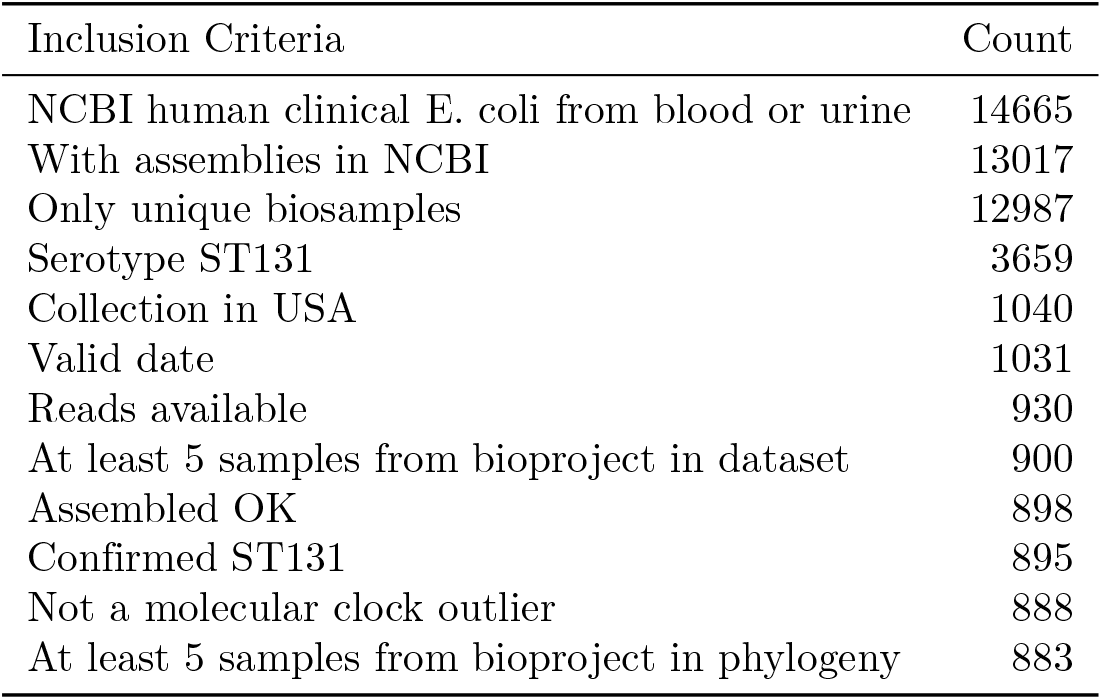

### S1 File Samples considered for analysis

Table with all ST131 clinical blood or urine samples from the USA considered for analysis. Each row lists the BioSample ID; whether it was included in the final analysis, and if not, gives the reason for exclusion; the BioProject from which it came; its associated assembly and isolate IDs; the IDs of any reads; and the location, specimen type, and collection date.

### S1 Fig. Feature correlation

Pairwise correlation between features colored by the relative magnitude of positive (red) or negative (blue) covariance.

### S2 Fig. Evolutionary history of *GyrA*

Visualization of ancestral reconstruction of *gyrA* presence (denoted by blue colored branches) or absence (gray branches). Trees are annotated with time and phylogenetic location of putative acquisition events (black) and loss events (red)

## Acknowledgments

This research is supported by the Centers for Disease Control and Prevention (CDC) (U01CK000587) and the US National Institutes of Health (NIH) (R35GM134934).

